# Y-chromosome haplotypes are associated with variation in size and age at maturity in male Chinook salmon

**DOI:** 10.1101/691063

**Authors:** Garrett J. McKinney, James E Seeb, Carita E. Pascal, Daniel E. Schindler, Sara E. Gilk-Baumer, Lisa W. Seeb

## Abstract

Variation in size and age at maturity is an important component of life history that is influenced both by environmental and genetic factors. In salmonids, large size confers a direct reproductive advantage through increased fecundity and egg quality in females, while larger males gain a reproductive advantage by monopolizing access to females. In addition, variation in size and age at maturity in males can be associated with different reproductive strategies; younger smaller males may gain reproductive success by sneaking in among mating pairs. In both sexes there is a trade-off between older age and increased reproductive success and increased risk of mortality by delaying reproduction. We identified four Y-chromosome haplogroups that showed regional and population-specific variation in frequency using RADseq data for 21 populations of Alaska Chinook salmon. We then characterized the range-wide distribution of these haplogroups using GT-seq assays. These haplogroups exhibited associations with size at maturity in multiple populations suggesting that the lack of recombination between X and Y-chromosomes has allowed Y-chromosome haplogroups to capture different alleles that influence size at maturity. Ultimately, conservation of life history diversity in Chinook salmon may require conservation of Y-chromosome haplotype diversity.

## Introduction

Variation in life history within populations is common across taxa and is often associated with alternative strategies for increasing fitness. This includes partial migration, where some individuals of a population migrate while others remain resident (Chapman et al. 2011), reproductive morphs that exhibit different mating strategies or sexually selected traits (Shuster 1989, Johnston et al. 2013, Küpper et al. 2015), age and size at maturity (Gibbons et al. 1981), and even length of life-span such as annual versus perennial plants (Hall et al. 2006). While life history variation is often assumed to be under the influence of many genes of small effect, examples of a large-effect genes and supergenes (sets of linked genes) influencing life history variation are increasingly being found. These include single genes that influence age at maturity (Barson et al. 2015) and sexually selected traits (Johnston et al. 2013); these also include chromosome inversions contributing to annual versus perennial life history (Twyford and Friedman 2015) and variation in migration versus residency (Pearse et al. 2014). The mechanisms underlying variation in life history have important implications for how this important diversity is maintained under different selective regimes.

Variation in life history strategies is exhibited by many salmon species with size and age at maturity being an important component of this variation. In females, large body size confers a direct reproductive advantage through increased fecundity and egg quality (Healey and Heard 1984). In contrast, variation in size and age at maturity in males can be associated with different reproductive strategies that exhibit frequency dependent fitness. For example, older larger males gain reproductive success by monopolizing access to females while younger smaller males gain reproductive success by sneaking in among mating pairs (Healey and Heard 1984, Berejikian et al. 2010). For both sexes, there is a trade-off to delayed maturation where increased reproductive success is tempered by an increased risk of mortality before reproduction. The optimal age at maturity for a population should represent the balance between reproductive benefits and mortality costs of delayed maturation (Healey 1986), and forces that change these costs or benefits could result in shifts in age composition. Nonetheless, considerable diversity in age at maturation is maintained within many populations, presumably as a bet-hedging strategy to spread risks over the life-cycle of these fish.

Age at maturity in salmon is generally thought to be a threshold trait that is dependent either upon reaching a minimum size at age or upon growth rate at key periods (Healey 1991, Thorpe 2007). Environmental factors that influence growth rate have been shown to influence age at maturity in many species. In the wild, studies show correlations between ocean conditions such as temperature and productivity and patterns of age at maturity (Otero et al. 2012, Siegel et al. 2017). In experimental settings, age at maturity was manipulated through changing temperature (Heath et al. 1994, Harstad et al. 2018) or food ration (Rowe and Thorpe 1990, Larsen et al. 2006).

In addition to environmental effects, multiple lines of evidence show a genetic component to age at maturity. High heritability values suggest considerable genetic variation for age at maturity in several salmon species (Gjerde 1984, Gall et al. 1988, Hankin et al. 1993, Heath et al. 1994), and Quantitative Trait Locus (QTL) and Genome-Wide Association (GWAS) studies identified genomic regions associated with variation in age at maturity (Barson et al. 2015, Kodama et al. 2018, Micheletti and Narum 2018, Waters et al. 2018). Studies also demonstrated that offspring of alternative male phenotypes exhibit different growth rates (Garant et al. 2002, Berejikian et al. 2011) with offspring of early maturation phenotypes (grilse and jacks) exhibiting high growth rates. In Atlantic salmon (*Salmo salar*), individuals with different life histories exhibit differing maturation thresholds that are genetically based (Aubin Horth and Dodson 2004). Despite these findings, the genetic basis of maturation age in most salmonids remains poorly understood.

One complicating factor is that the genetic architecture underlying variation in age at maturity appears to vary among salmonid species and even among populations within a species. In Atlantic salmon, age at maturity is strongly influenced by a single gene (VGLL3) (Ayllon et al. 2015, Barson et al. 2015). This gene exhibits sex-dependent dominance which facilitates sexually antagonistic selection (Barson et al. 2015). While this gene explained 39% of the phenotypic variability in European Atlantic salmon, studies in North American Atlantic salmon have shown that this association varies by population (Kusche et al. 2017, Boulding et al. 2019) and this gene has not been shown to have an effect in Pacific salmon (genus *Oncorhynchus*) (Micheletti and Narum 2018).

Chinook salmon (O. *tshawytschca* are the largest of the Pacific salmon and follow various life history strategies, spending 0 to 2 years in fresh water and 1 to 4 or more years in the ocean (Riddell et al. 2018). Male Chinook salmon exhibit significant variation in size and age at maturity (Healey 1991) that is linked to differential reproductive tactics and is likely controlled by both environmental and genetic components (Berejikian et al. 2010, Young et al. 2013). Many populations throughout North America recently experienced marked declines in size and age at maturity which may erode life history variation (Lewis et al. 2015, Ohlberger et al. 2018). Explanations for decreased age at maturity have focused on the impacts of fisheries induced evolution (Hard et al. 2008, Kendall et al. 2014) or changing environmental conditions (Siegel et al. 2017); however, these factors alone are not consistent or sufficient to explain current declines and it is likely that these declines are driven by multiple complex factors (Ohlberger et al. 2018). While the genetic control of age at maturity in Chinook salmon is still poorly understood, past studies offer clues to genomic regions that may be associated with maturation age. In particular, Heath et al. (2002) identified a strong sex-linked component to age at maturity in Chinook salmon, suggesting the influence of genes on the Y-chromosome.

The X and Y chromosomes in most salmonid species are morphologically undifferentiated (Davidson et al. 2009). Available sequence data suggests that the primary difference between sex chromosomes is an insertion containing the sex-determining gene (SDY, Yano et al. 2013), and in Chinook salmon a 2.4 Mb male-specific repetitive sequence has been identified (Devlin et al. 1998). While the sex-determining gene has been assigned to chromosome 17 (Ots17) in Chinook salmon (Phillips et al. 2013), this region is not in either of the current genome assemblies (Christensen et al. 2018, Narum et al. 2018) so the exact location of *sdY* is unknown.

Despite a lack of large-scale differentiation, the X- and Y-chromosomes could show sequence divergence due to sex-specific patterns of recombination. Recombination in females takes place along the full length of the chromosome while recombination in males is strongly localized to telomeric regions (Lien et al. 2011), restricting recombination between the X- and Y-chromosomes. Reduced recombination between sex chromosomes is supported by a 33Mb signal of sex association observed in Atlantic salmon (Kijas et al. 2018) and a lack of recombination between the sex-determining region and an allozyme locus on the sex-chromosome in Chinook salmon (Marshall et al. 2004). In addition to facilitating divergence between sex chromosomes, sex-limited recombination could lead to the formation of different Y-chromosome haplotypes and the capture of adaptive genetic variants. If Y-chromosome haplotypes have sufficiently diverged from the X-chromosome, they may be identified through patterns of extended linkage disequilibrium.

We examined patterns of linkage disequilibrium on the sex chromosome of Chinook salmon to determine if male-specific haplotype blocks (Y-chromosome haplotypes) existed, and if so, are these haplotypes associated with variation size and age at maturity which commonly differ between sexes in Chinook salmon. We identified four Y-chromosome haplogroups in Chinook salmon from Alaska that showed regional and population-specific variation in frequency. These haplogroups showed associations with size at maturity in multiple populations suggesting that the lack of recombination between X and Y-chromosomes has allowed Y-chromosome haplogroups to capture different genetic variants influencing size and age at maturity.

## Materials and Methods

### RAD Y-chromosome Haplotypes

We used existing RADseq data to examine patterns of genetic variation in the sex chromosome. RADseq data for 21 populations of Alaska Chinook salmon were obtained from: NCBI SRA accessions SRP034950 (Larson et al. 2014) and SRP129894 (McKinney et al. 2018), bioproject PRJNA560365 (McKinney et al. 2020b), and raw data used in (Dann et al. 2018). Raw data from (Dann et al. 2018) are available from those authors upon request. Locations of populations ranged from Cook Inlet to the Upper Yukon River and include a total of 1,082 samples (Figure 1, Table 1). RADseq data were processed with Stacks V1.7 (Catchen et al. 2011, Catchen et al. 2013) using default settings with the following exceptions: process_radtags (-c -r -q -filter_illumina -t 94), ustacks (-m 2 -M 2, -model_type bounded -- bound_high 0.05), cstacks (-n 2). The catalog of variation from McKinney et al. (2020b) was used for consistent RADtag names among this and previous studies. A total of six individuals per population from Cook Inlet was added to this catalog to allow for additional allelic variation.

**Figure 1.**
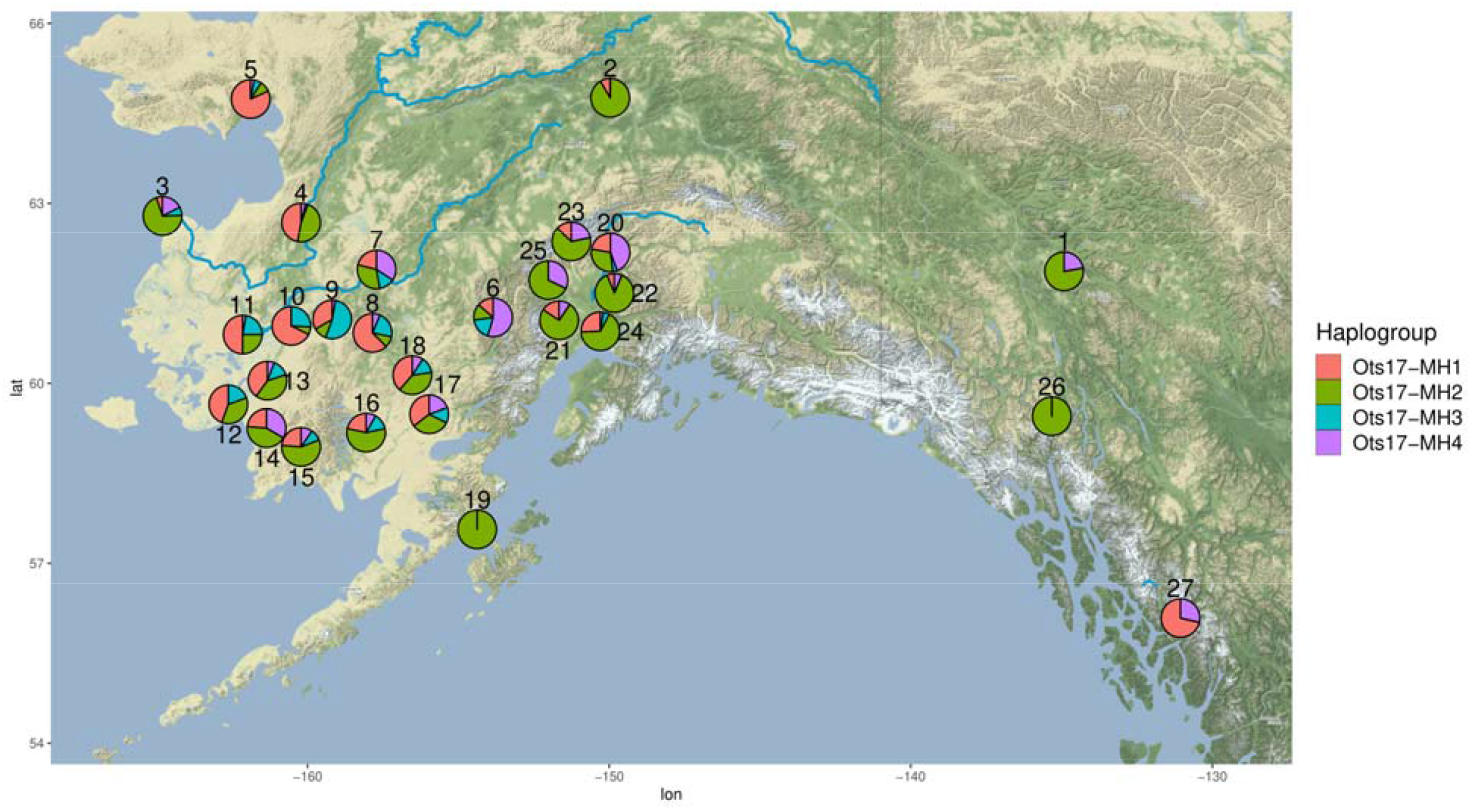
Frequency of Y-chromosome haplogroups throughout Alaska based on RADseq and GT-seq data. Locations of populations are approximate to prevent overlap of pie charts. Population names are given in Table 1. Note that the location for the Lower Yukon Test Fishery (population 3) indicates where fish were caught on their return to the Yukon River; these samples may represent fish from many populations that spawn throughout the Yukon River.

**Table 1.**
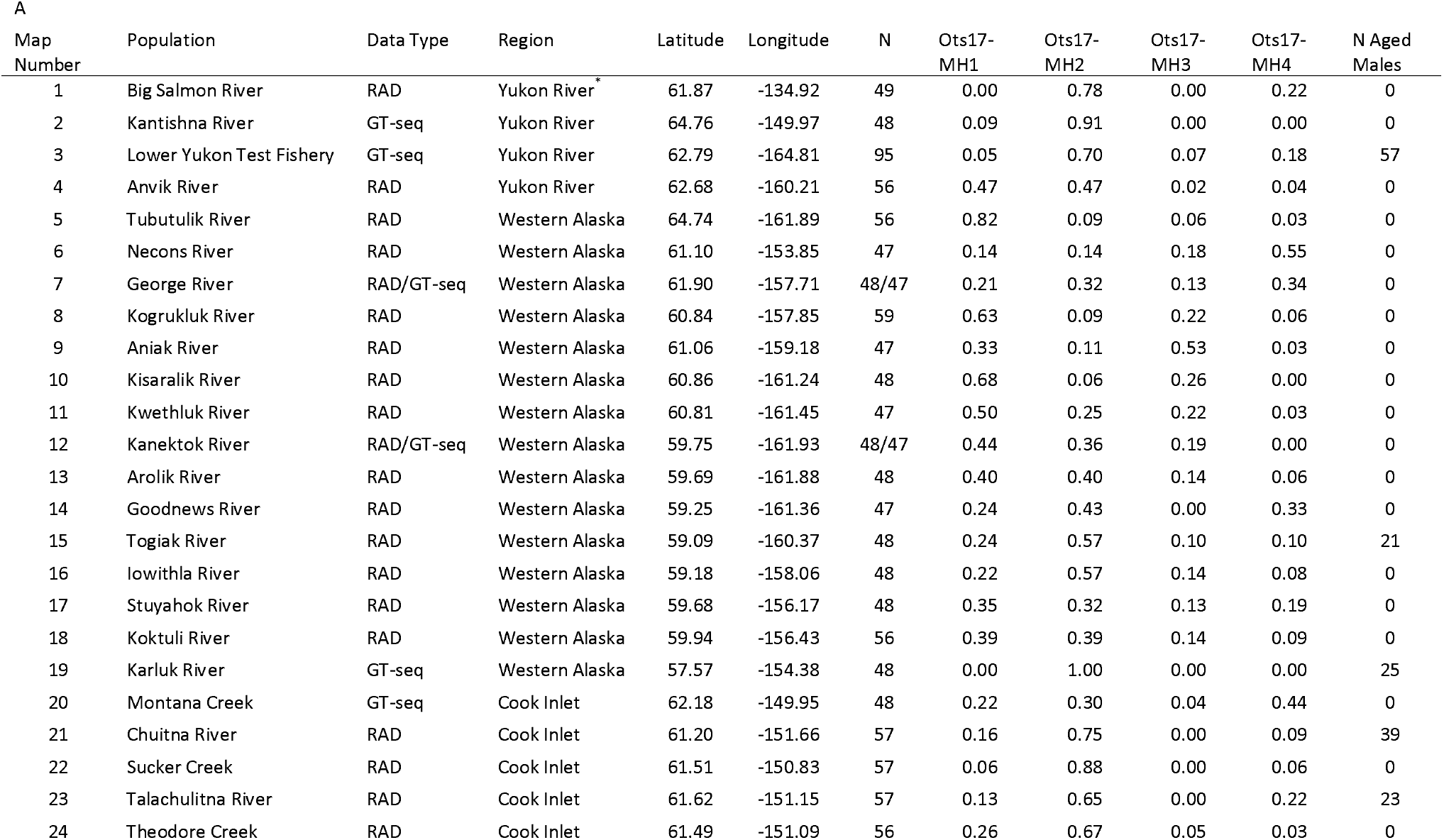
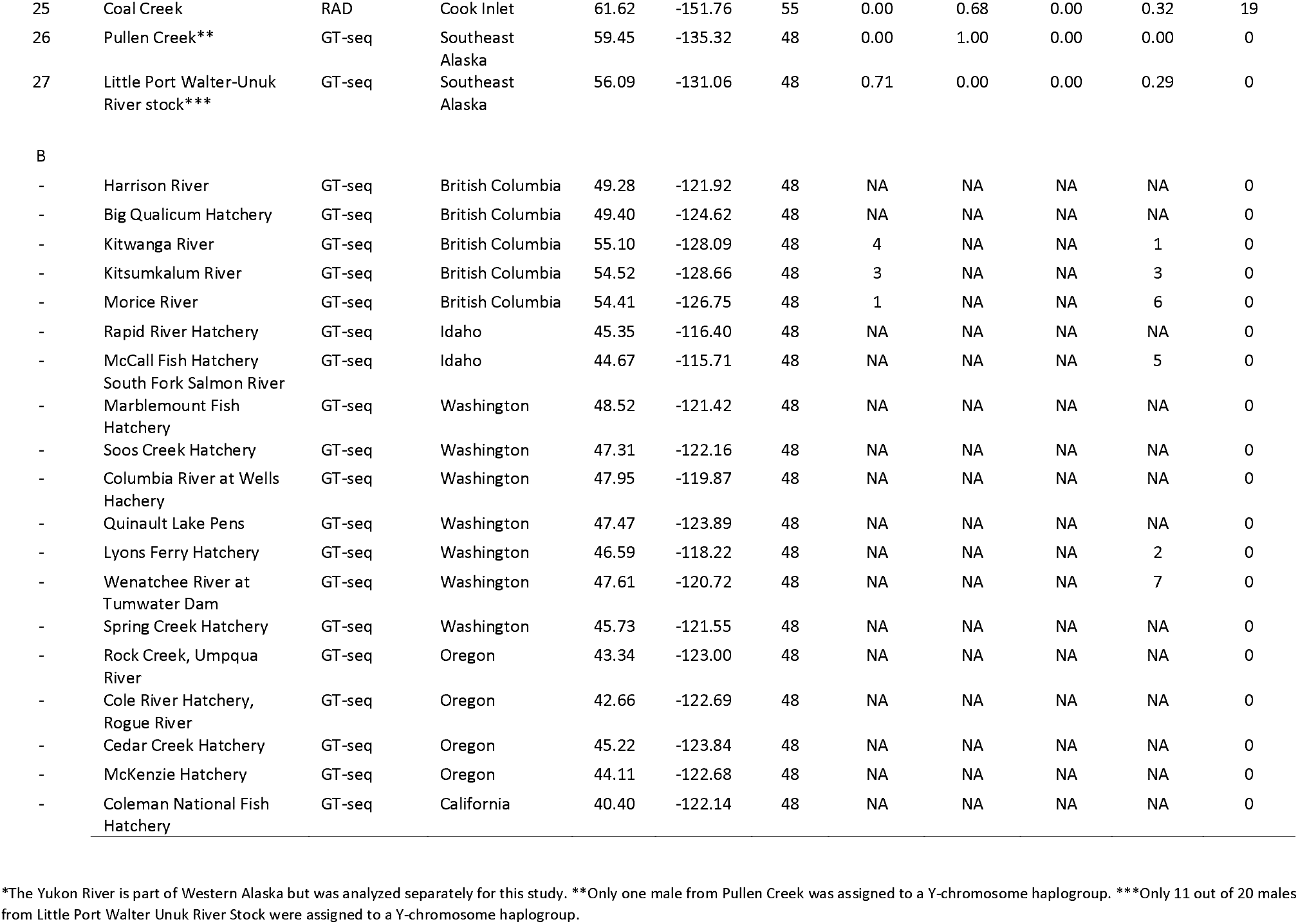
Populations used in this study. RADseq data was used for haplogroup discovery while GT-seq data was used to confirm that haplogroups were male-specific and to expand the geographic distribution of haplogroups. For populations with both RADseq and GT-seq samples, samples sizes for RADseq are given first. Frequency of Y-chromosome haplogroups are given for populations in Alaska in A. Due to the low proportion of males assigned to Y-chromosome haplogroups outside of Alaska, the number of individuals assigned to each Y-chromosome haplogroup for these populations are given in B.

RADseq results were filtered to remove SNPs that were likely to genotype poorly or be uninformative including collapsed paralogs, those with high missing data, and SNPs with low minor allele frequency. Paralogs are common in salmonid genomes due to an ancestral whole-genome duplication (Allendorf and Thorgaard 1984). With short read sequence the two copies of paralog are often collapsed into a single locus that appears polyploid; these cannot be reliably genotyped in typical RADseq studies because of insufficient read depth (McKinney et al. 2018). Collapsed paralogs were identified using *HDplot* (McKinney et al. 2017) and removed from further analysis. Loci with more than 10% missing data or a minor allele frequency (MAF) < 0.01 were also removed from analysis. Finally, loci were aligned to the Chinook salmon genome (Christensen et al. 2018) using Bowtie2 (Langmead and Salzberg 2012) to determine genomic position; only loci that aligned to the sex chromosome (Ots17, Phillips et al. 2013) with fewer than four mismatches were retained for analysis.

We identified putative Y-chromosome haplotypes by examining patterns of linkage disequilibrium on Ots17 using network analysis. We then phased genotypes for sets of high LD loci into haplotypes representing the alleles on each chromosome pair within an individual and clustered these haplotypes based on similarity into haplogroups which are groups of similar haplotypes. Network analysis of linkage disequilibrium is an effective approach for identifying genomic structures that exhibit high LD (Kemppainen et al. 2015) and had been used to identify distinct but overlapping haplotype blocks in chum salmon (McKinney et al. 2020a). Pairwise linkage disequilibrium was calculated using the *r*^2^ method in *Plink* (V1.9) (Purcell et al. 2007, Chang et al. 2015) and SNP pairs with *r*^2^ ≥ 0.3 were retained for network analysis. Network analysis was conducted using the igraph package in R (https://igraph.org/r) to identify sets of loci with high LD. Sets of loci that contained at least five SNPs and spanned at least 1 Mb were retained for further analysis. Genotypes for retained SNPs were then phased into haplotypes using fastPhase (Scheet and Stephens 2006). Putative Y-chromosome haplogroups were identified by clustering haplotypes using heatmap2 in R (Warnes et al. 2020) with the Ward.D clustering algorithm to minimize within group variance. Haplogroups that appear to be male-specific are hereafter referred to as Y-chromosome haplogroups and assigned names based on the following convention: chromosome number, MH for male haplogroup, followed by a sequential number, e.g. Y-chromosome haplogroups on chromosome 17 would be Ots17-MH1, Ots17-MH2, and so on.

Allele frequencies within each haplogroup were visualized using logo plots with the ggseqlogo package in R (Wagih 2017).

Assays for SNPs that were diagnostic for Y-chromosome haplotypes were assembled into an amplicon panel (GT-seq, Campbell et al. 2015) for expanded genotyping (see methods in McKinney et al. 2020b). Primers were designed using batch primer3 (You et al. 2008). Primer design used the consensus RAD sequence for each RADtag unless SNPs occurred within 20 bp of the end of the RADtag. In these cases the consensus sequences were aligned to the genome using bowtie2 (Langmead and Salzberg 2012), and genomic sequence flanking the RADtag was added to the consensus sequence for primer design. Panel optimization followed the methods of McKinney et al. (2020b); one round of sequencing using 80 individuals was conducted to identify loci that over-amplified or produced unreliable genotypes. Amplification levels and genotyping accuracy were assessed using GT-score (McKinney et al. 2020b).

### Expanded genotyping

Additional samples ranging from Alaska to California were genotyped using the GT-seq panel to establish the geographic distribution of the haplogroups (Table 1). In addition to the Y-haplogroup markers, this panel included the sex identification marker Ots_sexy3-1 (Hess et al. 2016a) to confirm that Y-chromosome haplotypes were present only in male fish. A total of 1,341 samples from 27 populations were genotyped (Table 1); phenotypic sex was known for 193 of these. A total of 94 RADseq samples were included in the GT-seq genotyping to examine concordance between the two datasets. Samples were sequenced on a HiSeq 4000, and data were processed and genotyped using GT-score (McKinney et al. 2020b) available at https://github.com/gjmckinney/GTscore.

Two methods were used to assign GT-seq samples to Y-chromosome haplogroups. First, samples were assigned Y-chromosome haplogroups using the same methods as the RADseq samples: phasing genotypes into haplotypes using fastPhase followed by haplotype clustering using heatmap2. Genotypes from RADseq samples were included in the GT-seq haplotype assignment to assess concordance between the discriminatory ability of the full RADseq marker set and the subset that that were successfully developed into GTseq markers. Second, the expected genotype patterns for males with each haplogroup were constructed assuming fixation of alleles on the X-chromosome and the Y-chromosome. The observed genotypes were then compared to the expected genotypes. Samples were assigned to a haplogroup if the observed genotypes had less than two mismatches to the expected genotypes.

### Y-chromosome haplotype analyses

Assignments were combined for GT-seq and RADseq samples to characterize the distribution and frequency of Y-chromosome haplogroups. Data for length and age at maturity were obtained from Alaska Department of Fish and Game (ADFG) for a subset of populations in Alaska and compared with Y-chromosome haplogroup data to determine if there were relationships between Y-chromosome haplogroup and length and age at maturity. Analysis of variance (ANOVA) was used to test the significance of associations between Y-chromosome haplogroups and length; population of origin was included as a covariate. A post-hoc Tukey test was then performed to determine if differences in size distribution were significant between individual haplogroups. The significance of associations between Y-chromosome haplogroups and age at maturity in the Lower Yukon Test Fishery were assessed using an ANOVA with population of origin added as a covariate. There are multiple methods of reporting age in salmon (Koo 1962); we report freshwater and ocean age for each individual using European notation, so an individual with an age of 1.3 would have spent 1 year growing in freshwater after emergence followed by 3 years in the ocean for a total age of 5 years.

## Results

A total of 448 SNPs remained after filtering SNPs with more than 10% missing data or a MAF ≤ 0.01, removing paralogs identified by HDplot, and retaining only SNPs that aligned to chromosome 17. Retained SNPs had an average coverage of 24x.

### RADseq Y-haplotype discovery

Network analysis of linkage disequilibrium identified three sets of linked SNPs exhibiting long-distance LD (4.5-9.5 Mb each). There were 35 SNPs in these high LD sets, representing 7.8% of the SNPs on Ots17 retained after filters. Individual genotypes for these SNPs were then phased, resulting in two haplotypes per individual. Clustering of haplotypes resulted in five major groupings that exhibited extended LD up to 17.4 Mb (Figure 2). Although phenotypic sex was available from only a subset of individuals, phased haplotypes from sexed individuals were present in each of these groupings (Table 2). One of the groupings contained phased haplotypes from both males and females suggesting that this represents the X-chromosome (gray cluster, Figure 2). Four of the haplogroups contained phased haplotypes predominantly from male samples (85%-95%), suggesting that these are from the Y-chromosome (Table 2, Figure 2). In addition, all individuals with a haplotype in one of the Y-chromosome haplogroups had their second haplotype in one of the X-chromosome groupings. All Y-chromosome haplogroups were present, at varying frequency, in each geographic region, suggesting that these haplogroups are conserved throughout this broad geographic region (Figure 1, Table 1, Table 3). Within Alaska, Y-chromosome haplogroups showed regional variation in frequency with the Ots17-MH1 haplotype exhibiting the high frequencies in most parts of Western Alaska and the Ots17-MH4 haplotype the most frequent in Cook Inlet (Figure 1). While some females had haplotypes within the Y-chromosome haplogroups, individuals had been visually sexed based on external features which is known to have variable accuracy (Lozorie and McIntosh 2014).

**Figure 2.**
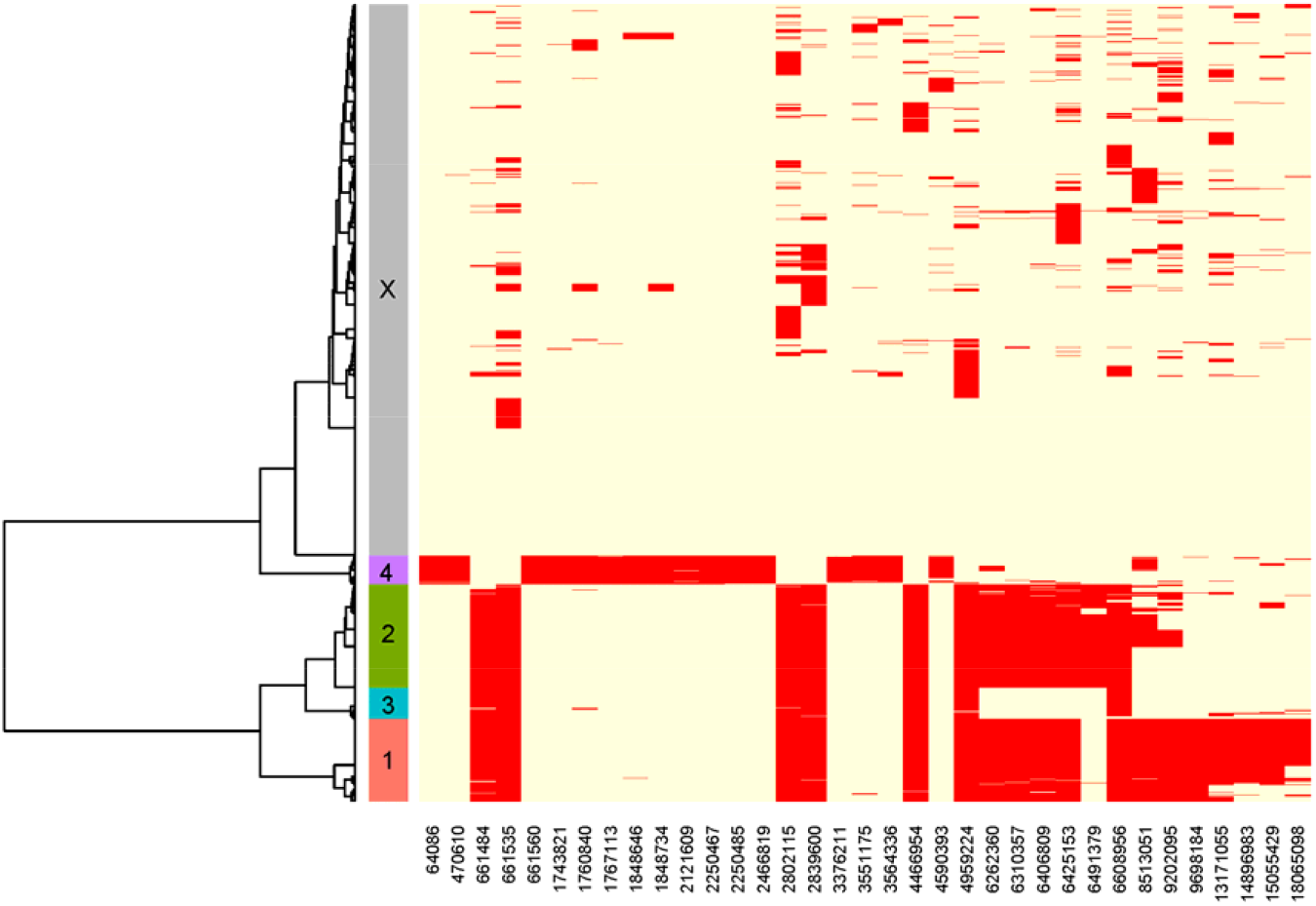
Plot of haplotype clusters identified by phasing high LD loci from the RADseq dataset. Each individual is represented by two haplotypes corresponding to each chromosome of Ots17. For each SIMP, the most common allele is in yellow and the least common allele is in red. Haplotypes were clustered into haplogroups, and six major haplotype clusters were identified; these are denoted by different colors along the sample dendrogram (y-axis). The gray haplogroup represents X-chromosomes while four haplogroups (pink=Ots17-MH1, green=Ots17-MH2, blue=Ots17-MH3, purple=Ots17-MH4) represent Y-chromosomes; numeric designations only are labeled on plot. SNP position on Ots17 is given on the x-axis.

**Table 2.**
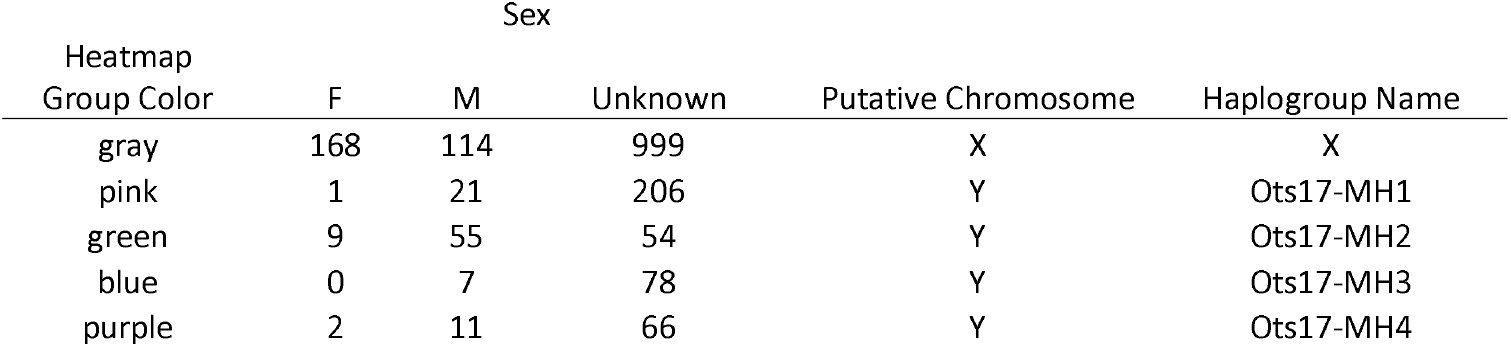
Number of haplotypes from RADseq samples assigned to each haplogroup by phenotypic sex. Each individual has two haplotypes. Males should have one haplotype assigned to a Y-chromosome haplogroup and one haplotype assigned to an X-chromosome haplogroup. Females should have both haplotypes assigned to an X-chromosome haplogroup.

| Heatmap<br>Group Color | Sex |  |  | Putative Chromosome | Haplogroup Name |
| --- | --- | --- | --- | --- | --- |
|  | F | M | Unknown |  |  |
| gray | 168 | 114 | 999 | X | X |
| pink | 1 | 21 | 206 | Y | Ots17-MH1 |
| green | 9 | 55 | 54 | Y | Ots17-MH2 |
| blue | 0 | 7 | 78 | Y | Ots17-MH3 |
| purple | 2 | 11 | 66 | Y | Ots17-MH4 |

**Table 3.** Number of haplotypes from RADseq samples assigned to each haplogroup by region. Each individual has two haplotypes. Males should have one haplotype assigned to a Y-chromosome haplogroup and one haplotype assigned to an X-chromosome haplogroup. Females should have both haplotypes assigned to an X-chromosome haplogroup.

| Heatmap<br>Group Color | Cook<br>Inlet | Western<br>Alaska | Yukon<br>River | Putative Chromosome | Haplogroup Name |
| --- | --- | --- | --- | --- | --- |
| gray | 410 | 942 | 139 | X | X |
| pink | 20 | 183 | 25 | Y | Ots17-MH1 |
| green | 112 | 130 | 39 | Y | Ots17-MH2 |
| blue | 2 | 82 | 1 | Y | Ots17-MH3 |
| purple | 20 | 53 | 6 | Y | Ots17-MH4 |

Allele frequencies within each haplogroup were visualized using logo plots (Figure 3). A total of 35 SNPs was found to have fixed or nearly-fixed differences in allele frequencies between Y-chromosome haplogroups and the X-chromosome. The Ots17-MH1, Ots17-MH2, and Ots17-MH3 haplogroups shared seven of the SNPs that differentiated these haplogroups from the X-chromosome (27492_30, 42530_30, 57320_53,64766_37,67724_90,5884_57,94509_60), suggesting a common evolutionary origin. The Ots17-MH4 haplogroup shared no diagnostic SNPs with the other Y-chromosome haplogroups. Several of the SNPs had allelic variation within the X-chromosome, (e.g., SNPs 57320_53 and 64766_37, Figure 3) suggesting that in some cases the Y-chromosome haplogroups have partitioned existing variation on the X-chromosome. Alternatively, some SNPs exhibited no allelic variation within the X-chromosome suggesting that alternative alleles observed on the Y-chromosome haplotypes may be novel genetic variants (e.g. SNP 71572_67, Figure 3).

**Figure 3.**
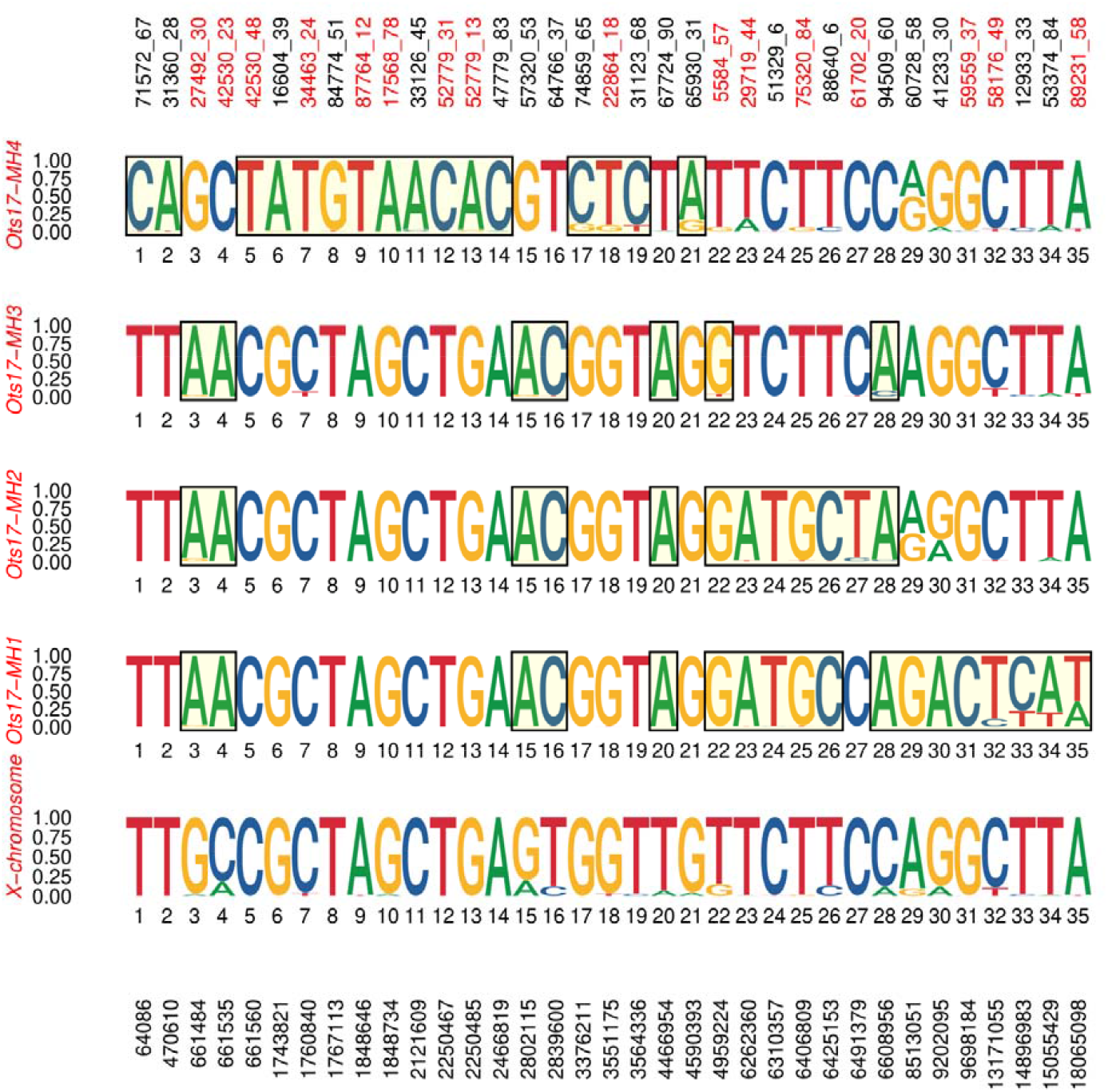
Logo plots showing allele frequencies within each of the male haplogroups and the X-chromosome for the RADseq samples. For each SNP, the frequency of alleles within each haplotype (range = 0-1) is shown by the height of the allele. Alleles which are putatively fixed relative to the X-chromosome are bounded by boxes. SNPs identified by tag number that were successfully converted to GT-seq assays are in red on the top X-axis. SNP positions on Ots17 are on the bottom X-axis.

### GT-seq Y-chromosome expanded sampling

A GT-seq panel was developed to genotype Y-chromosome haplotype markers for a set of samples representing the North American range of Chinook salmon. A total of 23 of the 35 RAD SNPs passed filtering criteria prior to primer design; of these 16 RAD SNPs were successfully converted to GT-seq assays (Figure 3, Table S1). Samples genotyped with GT-seq were assigned haplogroups using two methods, clustering of haplotypes using the Ward.D algorithm and assignment based on genotypes. Haplogroups Ots17-MH1 and Ots17-MH2 differed by several fixed SNPs in the RADseq dataset but only two fixed SNPs in the GT-seq panel (Figure 3). For these haplogroups an additional requirement was set that individuals must have the appropriate alleles for each of these fixed SNPs. For example, individuals in the Ots17-MH2 haplogroup must have a T allele at position 6491379 and a G allele at position 9698184.

Samples that were genotyped with both RADseq and GT-seq showed high concordance between RADseq and both GT-seq haplogroup assignment methods except for Ots17-MH3 (Table S2). When RADseq haplogroup assignment was compared with GT-seq haplogroup assignment using Ward.D clustering, 10 of the 12 samples assigned to RADseq Ots17-MH3 were grouped with females. This is likely due to the fact that only three of the RADseq SNPs that differentiated this haplotype from the X-chromosome were successfully developed into GT-seq (Figure 3). In contrast, when GTseq samples were assigned to haplogroups based on comparing observed to expected genotypes for each haplogroup, 10 of the 12 samples correctly assigned to Ots17-MH3 while two were not assigned to any haplogroup. For samples genotyped only with GT-seq, there were no discrepancies in haplogroup assignment between the clustering and genotype-based methods except that Ots17-MH3. Ots17-MH3 had no individuals assigned using haplotype clustering but had 23 samples assigned based on genotype matching. Genotype-based matching assigned approximately 12% fewer individuals to haplotypes overall than the phased haplotype clustering (Table S2C) but was better able to assign individuals to the Ots17-MH3 haplogroup. All GT-seq samples assigned to a Y-chromosome haplogroup were genetic males based on the Ots-SEXY-3-1 sex-identification assay.

The majority of male (phenotypic or genetic) Chinook salmon in Alaska were assigned to Y-chromosome haplogroups with regional variation in assignment rate. Overall, 90% of phenotypic males (93/103) and 87% of genetic males (232/266) were assigned to a haplogroup. Genetic males that were not assigned Y-chromosome haplotypes were concentrated in Southeast Alaska; 30 of the 40 unassigned males were from the Little Port Walter and Pullen Creek populations. Only 58% of males from Little Port Walter and a single male from Pullen Creek could be assigned to a haplogroup. These samples had low missing data suggesting that the low assignment rate is due to low prevalence of these haplogroups in this region. Excluding these two populations results in 98% haplogroup assignment of male Chinook salmon in Alaska. Distribution of Y-chromosome haplotypes varied regionally: Y-chromosome haplotype blocks identified in Alaska were present in nearly all genetic males within Alaska but had rare occurrence outside of Alaska (Table 1). In total, eight males assigned to Ots17-MH1 were found in British Columbia, and 24 males assigned to Ots17-MH4 were found in British Columbia, Washington, and Idaho.

### Size and Age at Maturity

Size at maturity data were available for nine populations in this study; five populations were genotyped using RADseq data, three were genotyped using GT-seq data, and one was genotyped using both RADseq and GT-seq. Populations were grouped by region (Yukon River, Western Alaska, and Cook Inlet) for visualization. Boxplots of size at maturity for each Y-chromosome haplotype showed a consistent relationship throughout these regions (Figure 4). The Ots17-MH1 haplogroup had the smallest individuals, the OTS17-MH2 haplogroup was associated with intermediate sized fish, and the Ots17-MH4 haplogroup was associated with the largest individuals in each region. The Ots17-MH3 haplogroup showed inconsistent results with small fish in some regions and large fish in others. ANOVA results showed that both haplogroup and population were significantly associated with variation in size at maturity. Post-hoc Tukey tests were conducted to determine which haplotypes had statistically significant differences in size. In the Yukon River, individuals with the Ots17-MH4 haplotype were significantly larger (p<0.05) than fish with the Ots17-MH1 or Ots17-MH2 haplotype. While this pattern was repeated in Western Alaska, the relationship was not statistically significant, likely due to low sample size of the Ots17-MH4 haplogroup (Figure 4). In Cook Inlet, the Ots17-MH1, Ots17-MH2, and Ots17-MH3 all had significant differences in size.

**Figure 4.**
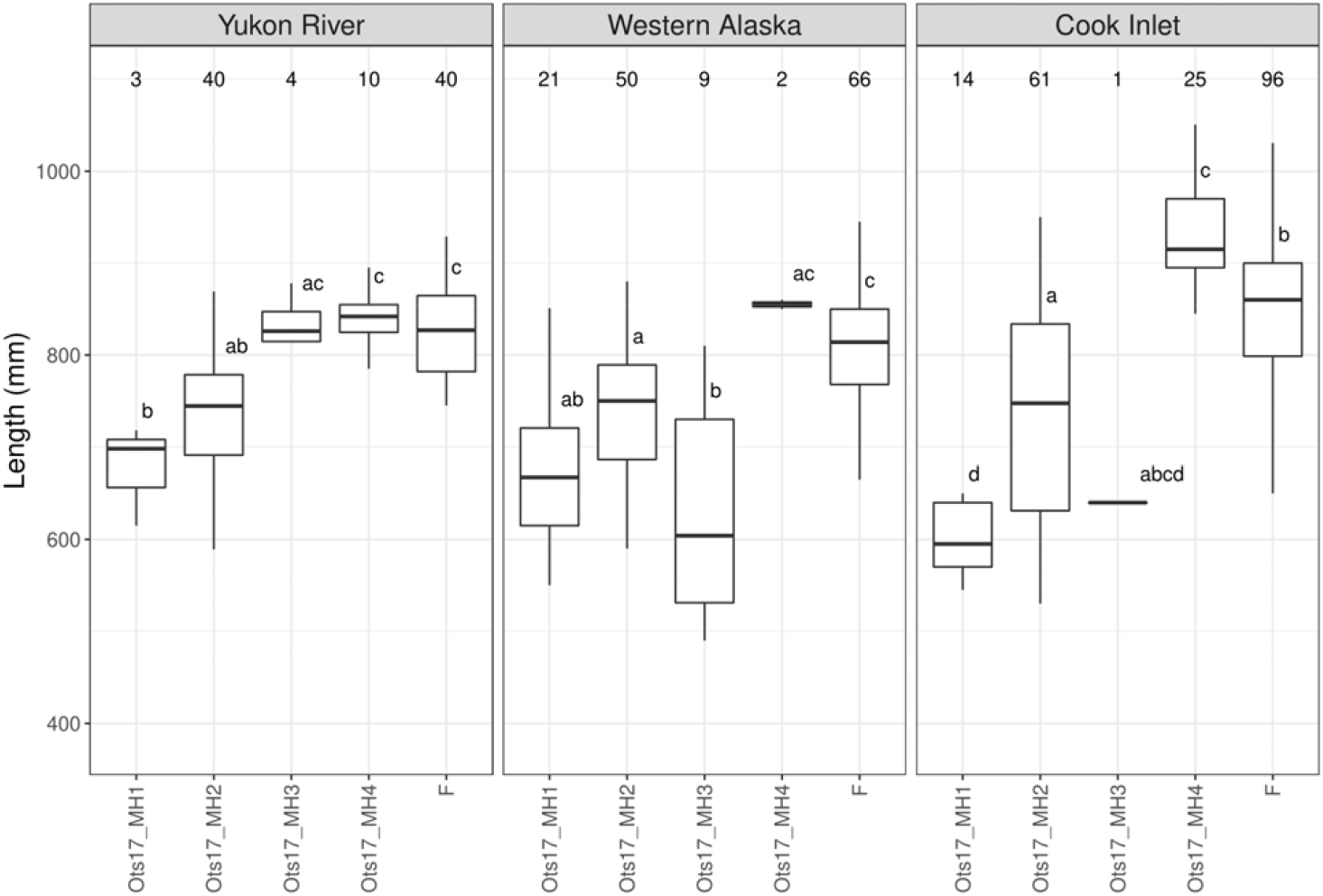
Distribution of size at maturity for each Y-chromosome haplogroup for Alaska Chinook salmon. Female (F) size at maturity is included for comparison. Samples sizes for each haplogroup are given above the boxplots. Within each region, Y-chromosome haplogroups with statistically different lengths are represented by different letters. Haplogroups with two letters (i.e., ab) do not have statistically different size distributions from haplogroups with a or b.

### Age at maturity data were available for 177 males from six of the populations in this study

(Table 1). The number of males with age data in each population ranged from 19 (Coal Creek) to 57 (Lower Yukon Test Fishery). There was a significant association between Y-chromosome haplogroup and age at maturity (p=0.047). The distribution of age at maturity by haplogroup was plotted for the Lower Yukon Test Fishery as this collection had the most samples with age data. Histograms of age at maturity for each Y-chromosome haplotype revealed that the Ots17-MH2 haplotype had approximately three times as many 1.3 fish as 1.4 fish while the Ots17-MH4 haplotype had nearly even proportions of 1.3 and 1.4 fish (Figure 5). Boxplots of size at age showed that fish with the Ots17-MH4 haplotype were larger for age 1.3 and 1.4 than fish with the Ots17-MH2 haplotype. Results were statistically significant for age 1.3 fish but not age 1.4, possibly due to low sample size. The smallest fish had the Ots17-MH1 haplotype; however, only two fish in the Lower Yukon Test Fishery had both size and age data for this haplotype.

**Figure 5.**
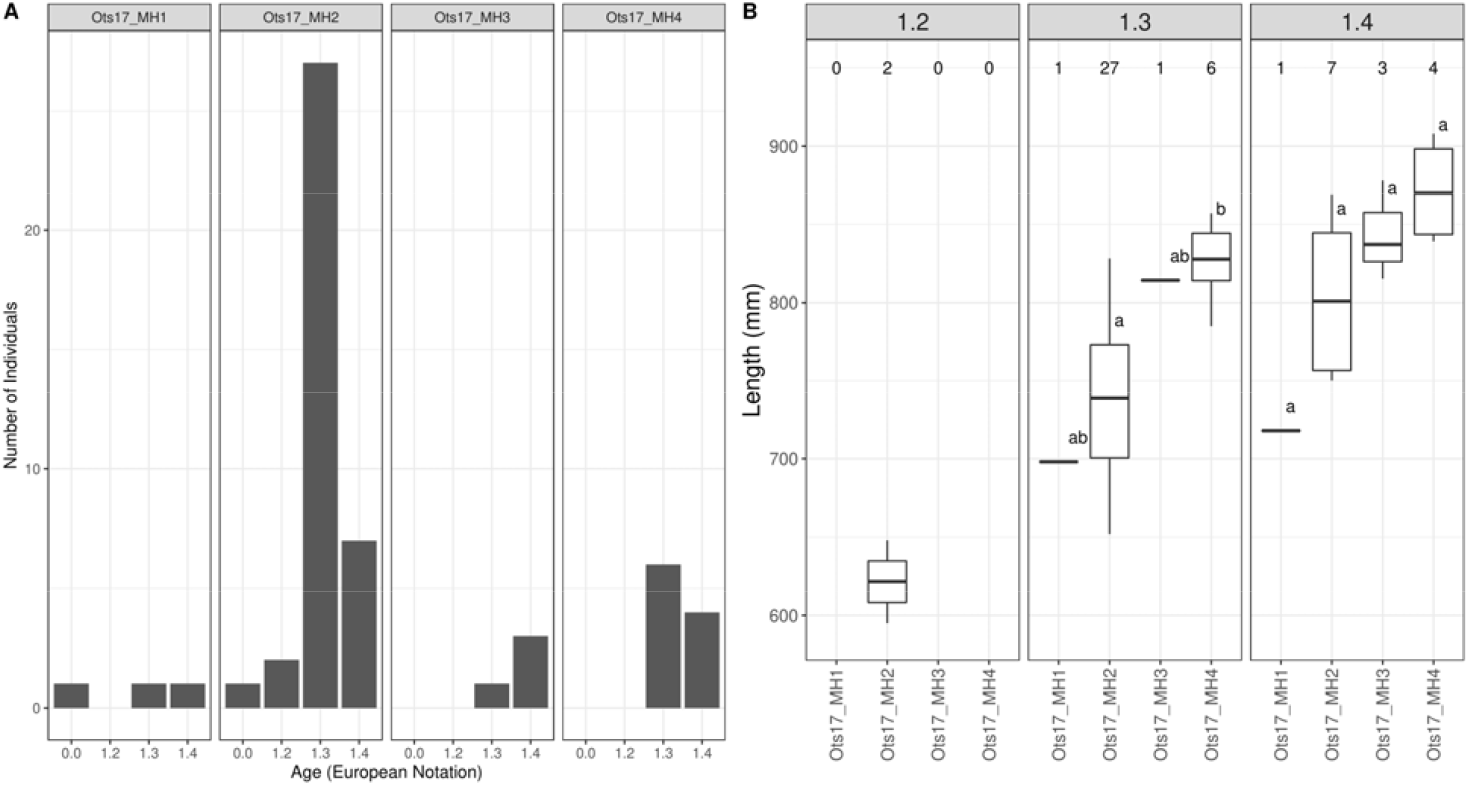
Distribution of A) age at maturity and B) size at age for each Y-chromosome haplogroup in the Yukon River. Within each age class, Y-chromosome haplogroups with statistically different lengths are represented by different letters. Sample sizes for each haplotype are given above the boxplots.

## Discussion

Variation in life history within populations is common across taxa and is often assumed to be under the influence of many genes of small effect; however, examples of a genes and regions of large effect on life history variation, including single genes, small genomic regions, or chromosome inversions, are increasingly being found (Johnston et al. 2013, Pearse et al. 2014, Barson et al. 2015, Twyford and Friedman 2015). The genetic mechanism underlying variation in life history has important implications for how life history diversity is maintained under different selective regimes (Hess et al. 2016b, Prince et al. 2017), particularly in the case of sexually antagonistic selection where males and females have different fitness optima (Barson et al. 2015, Pearse et al. 2019).

Size and age at maturity are ecologically and evolutionarily important traits in Chinook salmon. Numerous studies have examined ongoing declines in age at maturity; however, it has been difficult to disentangle the interactions of environmental and genetic causes of this decline. Size and age associated markers and genes have previously been identified in genetic studies of Chinook salmon (Micheletti and Narum 2018, Waters et al. 2018); however, results were not consistent across populations, and no markers were located on the sex chromosome. We show that a conserved set of Y-chromosome haplotypes is associated with variation in size and age at maturity in Chinook salmon across the Yukon River, Western Alaska, and Cook Inlet. These observations open a new line of research into the genetic basis of age at maturity in salmonids.

### Range-wide distribution of haplotypes

Chinook salmon are represented by multiple genetically distinct lineages throughout the species range (Waples et al. 2004, Beacham et al. 2006, Moran et al. 2012). These lineages often show little gene flow among them due to differences in geographic range or spawn timing, and this isolation may result in different sets of Y-chromosome haplogroups and patterns of recombination across lineages. The Y-chromosome haplogroups that we identified through extended linkage disequilibrium were consistently observed throughout Chinook salmon populations from the Upper Yukon River south to Little Port Walter in Southeast Alaska. The occurrences become rarer in Southeast Alaska, and few individuals south of Southeast Alaska could be assigned haplotypes suggesting that the haplotypes identified within Alaska are regionally restricted. This corresponds with observed breakpoints between Chinook salmon lineages near Cape Fairweather (Templin et al. 2011) which is approximately 320 km northwest of Little Port Walter and 160 km west of Pullen Creek. While rare, the Ots17-MH4 haplogroup was found in some individuals as far south as Idaho in the northwestern continental United States, suggesting that this may be broadly distributed even if at low frequency. It is likely that other Y-chromosome haplogroups exist outside of Alaska, but we were unable to identify them with this dataset. The expanded survey used only a subset of SNPs, rather than the full RADseq dataset, that characterized the Y-chromosome haplogroups in Alaska. If different SNPs are associated with Y-chromosome haplogroups in other regions of the species range, we would not be able to identify them in this study. The presence of additional haplogroups outside Alaska could be determined by examining reduced-representation or whole-genome sequence data from additional populations.

Populations throughout Alaska showed variation in haplotype frequency which may be ecologically significant given the association between haplotypes and size and age at maturity.

Populations from Western Alaska and the Anvik River in the Lower Yukon River had a greater proportion of the Ots17-MH1 haplogroup which was associated with smaller fish. Populations in the Middle and Upper Yukon River and Cook Inlet were primarily composed of Ots17-MH2 and Ots17-MH4 haplogroup which were associated with larger fish. While we did not have enough samples with size data to characterize size distributions within regions, this finding is consistent with a long-term analysis of Chinook salmon returns by Lewis et al. (2015) who reported smaller fish on average in Kuskokwim and Nushagak river populations from Western Alaska relative to Yukon River and Cook Inlet populations. In addition, the Cook Inlet populations sampled in this study are near and share a common migration pathway with the Kenai River which has historically produced large Chinook salmon (Lewis et al. 2015, Schoen et al. 2017). The relationship between size and haplotype also varied by region, with the Yukon River having the smallest difference in sizes between haplotypes and Cook Inlet having the largest difference in sizes (Figure 4). While the magnitude of difference appears to be largest in Cook Inlet, it is difficult to accurately assess statistical significance due to the low sample size when splitting samples among regions. Taken together, these results suggest that differing frequencies of Y-chromosome haplotypes may contribute to regional variation in size of Chinook salmon and that the effect of haplotype on size can vary between regions, potentially due to other genetic influence or different environmental conditions. The Ots17-MH3 haplogroup was unusual in that it showed no consistent pattern, with large fish in some regions and small fish in other regions (Figure 4). This haplogroup also showed the least differentiation from the X-chromosome based on RADseq data (Figure 3) and may not contain adaptive variants influencing size or age at maturity.

### Resolving sexual conflict

Different fitness optima for males and females are common across species and are difficult to resolve when adaptive loci are on autosomes. One mechanism is for the same alleles to exhibit sex-specific dominance, such as the VGLL3 gene which influences age at maturity in Atlantic salmon (Barson et al. 2015). Another mechanism is to partition adaptive variants between non-recombining regions of sex chromosomes, such as genes governing coloration and fin morphology in Poecilids; these genes are attractive in males but would increase predation risk in females (Anna Lindholm and Felix Breden 2002). The existence of Y-chromosome haplotypes demonstrates not only that genetic variation is partitioned between the X- and Y-chromosomes, but that Y-chromosome variants have partitioned different genetic variants (Figure 3). While it is unlikely that the specific SNPs we observed are adaptive, these haplotypes are associated with variation in adaptive traits. This suggests that different adaptive variants have been captured by Y-chromosome haplotypes. Y-chromosome haplotypes also varied in the chromosomal regions where the X- and Y-chromosome were differentiated. SNPs that differentiated the Ots17-MH1, Ots17-MH2, and Ots17-MH3 haplogroups from the X-chromosome were generally concentrated from approximately 4 Mb up to 22 Mb (Fig. 2, 3). One exception was a single SNP at ~600 Kb. SNPs that characterize the Ots17-MH4 haplogroup were concentrated from ~600 Kb to 6Mb along chromosome 17. The haplogroups may have captured different adaptive variants in each of these regions. This partitioning of genetic variation can resolve sexual conflict in age at maturity and provide a mechanism for the evolution of life history diversity in males.

### Importance of Y-haplotypes for life history diversity

Male Chinook salmon exhibit life history diversity related to maturation age. Older, larger males are believed to have greater reproductive success through their ability to monopolize access to females. Males that mature younger and smaller as jacks are generally believed to have reduced reproductive success but have greater survival to maturation due to reduced risk of ocean mortality (Berejikian et al. 2010). These alternate life histories of male Chinook salmon may exhibit frequency dependent fitness which in theory should exhibit stable proportions; however, this assumes populations are at equilibrium. Male sockeye salmon (*O. nerka*) exhibit similar frequency dependent life history variation, but persistent demographic shifts towards an abundance of jacks (males that mature one year earlier than the earliest maturing females) has occurred in some populations as a result of strong selection events coupled with variation in recruitment (DeFilippo et al. 2019). If maturation size and age in males are strongly influenced by an individual’s Y-chromosome haplotype, then size-selective fishing practices or even size selective predation by marine predators (Ohlberger et al. 2019, Seitz et al. 2019) may result in shifts in haplogroup frequency, demographic changes, and loss of age diversity that are difficult to recover. The markers we developed can be used to characterize historic (i.e., from archived samples) and current Y-chromosome haplogroup diversity in Alaska Chinook salmon to determine if demographic shifts correspond with shifts in frequencies of Y-chromosome haplogroups. Ultimately, conservation of life history diversity in Chinook salmon may require conservation of Y-chromosome haplogroup diversity.

Hypotheses of population structure and delineation of management units using genetic data are typically based on genome-wide analyses consistent with the assumption that major life history traits are controlled by many genes with small effects. Waples and Lindley (2018) recently commented on the new challenges facing existing conservation frameworks when associations are identified between one or a very few genes and key life history traits. Their comment was prompted by the recent identification of SNPs from a GREB1L gene that explain a large proportion of the variation associated with seasonal timing of adults returning to spawn steelhead (*O. mykiss*) and Chinook salmon (Hess et al. 2016b, Prince et al. 2017). Conservation of the Y-chromosome haplotype shares similar challenges to the GREB1L situation. Waples and Lindley (2018) pose a series of key questions to help provide an informed basis for decisions or management actions. Among other questions, they argue that a full understanding of the distribution of the variation in space and time is needed and that investigations into the genes and mechanisms responsible for the life history variation should be initiated. In the case of the Y-haplotypes, additional questions exist such as: What are the causal variants within these haplotype blocks? Do haplotype blocks exhibit consistent phenotypes in different environments and with different genetic backgrounds? What proportion of variance in size and age at maturity are explained by these haplotype blocks relative to other regions of the genome? If haplotype blocks are found in other regions of the Chinook salmon range, do they function in a similar manner to those suggested by the results of this study?

## Conclusion

Variation in size and age at maturity is common across taxa and is often associated with alternative strategies for increasing fitness; however, the genetic basis of this variation is largely unknown. We identified Y-chromosome haplogroups that are associated with size, and likely age, at maturity in Chinook salmon throughout Alaska. These haplogroups were primarily restricted to western and southcentral Alaska Chinook salmon where the most diversity in age-at-maturity exists, and likely represent a subset of the total diversity across the species range. Interestingly, this region holds the most stable populations of Chinook salmon in the Northeast Pacific (Griffiths et al. 2014), possibly due to the bet-hedging benefits of diversity in age at maturity. It is possible that each Chinook salmon lineage has a specific set of haplogroups and relationships between haplotypes and size/age at maturity may differ by lineage. The discovery of Y-chromosome haplotypes and their potential effect on life history variation in Chinook salmon may help explain the causes and consequences of the recent declines in size and age of adult Chinook salmon, trends that are most pronounced in the region with the highest haplotype diversity. Ongoing efforts to understand the causes of these declines point to size-specific mortality of maturing fish but also require an unknown evolutionary basis (Ohlberger et al. 2019). The results presented here point to such a mechanism for the genetic control of changes in size-at-age and age-at-maturity in Chinook salmon, as future changes in environmental conditions and selective fishing may lead to further demographic responses in this economically and ecologically important species.

## Acknowledgements

We would like to thank Chris Habicht, Bill Templin, Wes Larson, and Todd Seamons for helpful discussions and support for this research. This research was partially funded by North Pacific Research Board grant 1724-000, AKSSF grants 44812, 44913 and 44914, and Pacific States Marine Fisheries Commission grant # 16-108G to Alaska Department of Fish and Game. GJM was partially supported by a National Research Council postdoctoral fellowship. The statements, findings, and conclusions are those of the authors and do not necessarily reflect the views of the NOAA, the U.S. Department of Commerce, or the ADFG.

## Data Accessibility

GT-seq data will be deposited to NCBI SRA upon acceptance.

## Supporting Information

**Table S1:** Primer and probe sequence for GT-seq loci that were used for expanded genotyping of Y-chromosome haplotypes.

**Table S2:** Concordance between methods of Y-haplogroup assignment. Samples genotyped with both RADseq and GT-seq are shown in A and B. GT-seq samples were assigned to haplogroups using a clustering algorithm in A and by matching observed genotypes to expected reference genotypes in B. Samples genotyped only with GT-seq are shown in C.

